# TNF signaling drives expansion of Reg4+ epithelial cells in colitis

**DOI:** 10.1101/2025.09.08.674949

**Authors:** Safina Gadeock, Nandini Girish, Cambrian Y. Liu, Ying Huang, Tracy C. Grikscheit, D. Brent Polk

## Abstract

Reg4+ secretory cells are upregulated in colitis and promote epithelial repair, but their regulation is poorly understood. We show that TNF-TNFR1 signaling controls Reg4+ cell numbers in mouse and human colonoids and in vivo, with TNFR1 deficiency reducing these cells and TNF restoring them dose-dependently. In UC patients and DSS-colitis, REG4+ cells mark regenerating crypts, and persistent REG4, DUOX2, and TNFR1 expression identifies non-responders to anti-TNF therapy. These findings reveal a TNFR1-dependent mechanism regulating Reg4+ cells and suggest potential strategies to improve therapeutic responses in IBD.

## Introduction

Regenerating islet-derived (REG)4 marks a secretory cell subset in human and mouse colonic epithelium and is highly upregulated in IBD (1). In mice, Reg4+ deep crypt secretory (DCS) cells support regeneration, stem cell function (2), and host defense (3). However, whether Reg4 upregulation in colitis reflects DCS cell expansion, and if IBD-associated cytokines directly specify Reg4+ cell development, remains unclear. Here, we identify a novel mechanism by which TNF regulates Reg4+ secretory cell numbers via TNFR1 signaling in human and mouse colitis, shedding light on TNF’s role in epithelial remodeling in IBD.

## Results

To better define REG4^+^ cells in the healthy human and mouse colon, we first examined their crypt localization. In control (non-IBD) pediatric patients, REG4 mRNA and protein localize to discrete mucous cells in the differentiated upper half of the colonic crypt (Fig. 1*A*), contrasting with the crypt base localization of Reg4 in the healthy mouse colon. In pediatric colitis patients, REG4 expression increases overall (Fig. 1*B*), extending from surface epithelial cells to discrete cells at the crypt base (Fig. 1*A*), resembling Reg4^+^ localization in DSS-induced colitis (Fig. 1*C*). These findings indicate that REG4 is not a marker of a DCS lineage in the healthy human colon, but that colitis drives a conserved remodeling in both species, expanding REG4 expression within subsets of secretory epithelial cells. TNF-TNFR1 signaling has been implicated in regulating intestinal epithelial differentiation (4, 5). To test its role in Reg4+ DCS cell expression, we used TNFR1^ΔIEC^ mice lacking epithelial TNFR1. These mice showed a 36% reduction in Reg4+ cells and a 2.7-fold decrease in Reg4 mRNA compared to TNFR1^fl/fl^ controls (Fig. 1*D*). Notably, while the Reg4+ zone was reduced, crypt height remained unchanged (Fig. 1*E*), suggesting compensatory mechanisms during normal cell turnover. Similarly, TNFR1^-/-^ colonoids showed a 2.3-fold decrease in Reg4 mRNA and a 55% drop in Reg4+ cells versus TNFR1^+/-^ controls (Fig. 1*F*), supporting an epithelial-intrinsic role for TNFR1 in maintaining homeostatic Reg4+ cell expression.

**Fig. 1.**
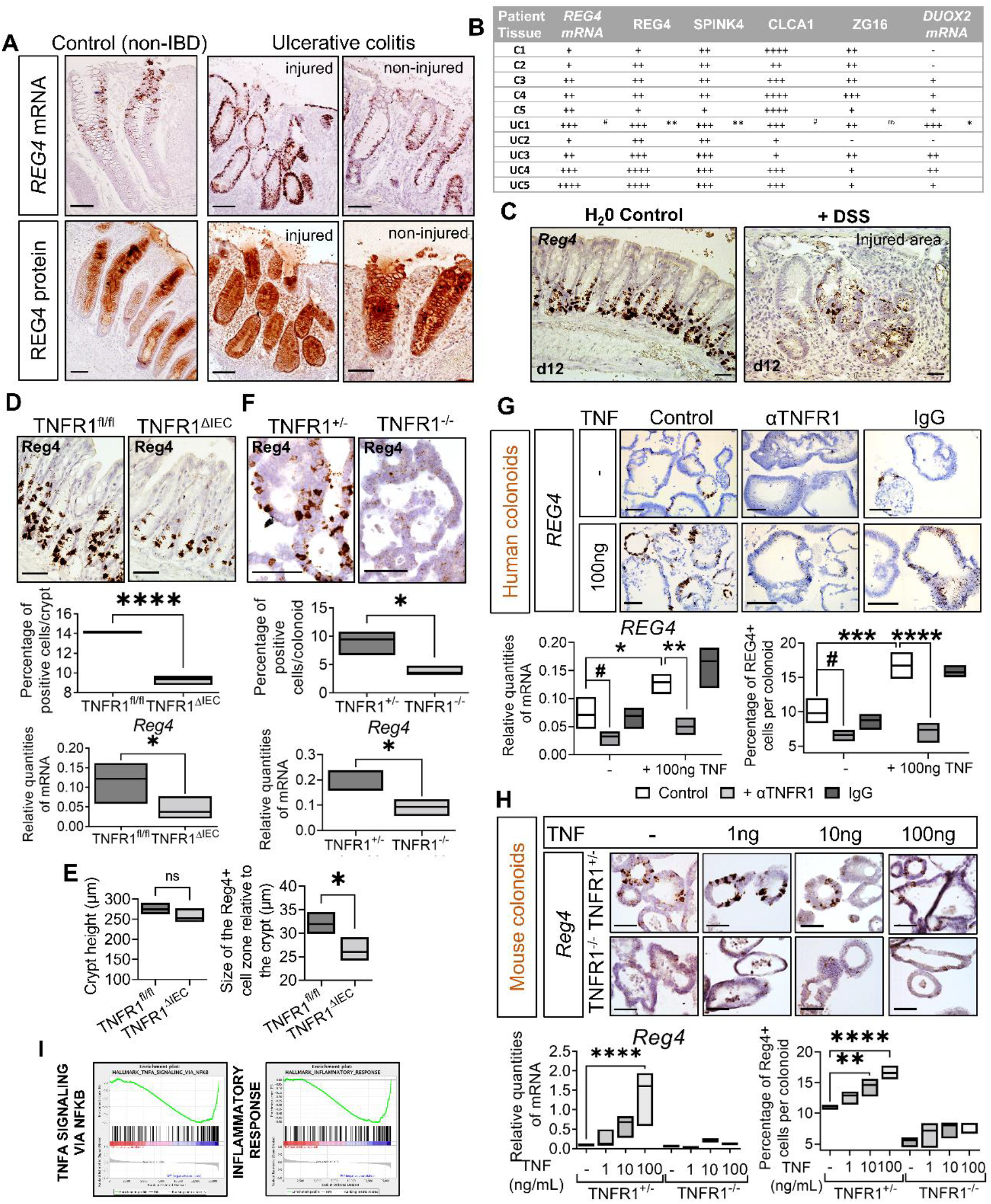
REG4 localized by ISH and IHC in pediatric non-IBD and UC colon (A). REG4, SPINK4, CLCA1, ZG16, and DUOX2 expression was quantified using Human Protein Atlas scoring; t-test, *p < 0.05, **p < 0.01, ^#^p = 0.05–0.09 (n = 5) (B). REG4 localized to UC surface epithelium and Reg4 to DSS-treated mouse colon (2% DSS, 6 d + 7 d recovery; n = 3-5) (C). Reg4 ISH, Reg4^+^ cell counts, and qRT-PCR in TNFR1^fl/fl^ and TNFR1^ΔIEC^ mice colon (n = 3-5) (D), and crypt height/Reg4^+^ zone length (E). Reg4 ISH, Reg4^+^ cell counts, and qRT-PCR in TNFR1^+/−^ and TNFR1^−/−^ colonoids (n = 3) (F). REG4 ISH, qRT-PCR, and % REG4^+^ cells in human colonoids treated with TNF ± anti-TNFR1 antibody or IgG (n = 3) (G). Reg4 ISH, qRT-PCR, and % Reg4^+^ cells in TNFR1^+/−^ and TNFR1^−/−^ colonoids treated with TNF (n = 3) (H). GSEA shows TNF, IFN-γ, and inflammatory pathway enrichment in TNFR1^+/−^ vs TNFR1^−/−^ colonoids (n = 3) (I). G–H analyzed by one-way ANOVA with Tukey’s test; ^#^p = 0.05–0.09, *p < 0.05, **p < 0.01, ***p < 0.001, ****p < 0.0001.

While TNF’s role in regulating mucus production is well established, its influence on epithelial secretory cell fate in colitis has mostly been explored in injury models (4) or via genetic TNF manipulation (5). To investigate TNF’s role in Reg4^+^ cell expression, we used human and mouse colonic organoids. Continuous TNF treatment significantly increased REG4^+^ cell numbers and expression in human colonic organoids from non-IBD controls, an effect blocked by anti-TNFR1 neutralizing antibody (Fig. 1*G*). Similarly, TNF promoted dose-dependent recovery of Reg4^+^ cells in mouse organoids, which was abolished in TNFR1^−/−^ organoids (Fig. 1*H*). Notably, organoids without exogenous TNF still showed baseline TNFR1-mediated signaling, likely from epithelial-derived TNF (Fig. 1*I*). Thus, Reg4^+^ cells are TNF-responsive in human and mouse colonic epithelia.

To assess whether inflammation drives Reg4^+^ cell expansion in vivo, as seen in human UC, we examined TNFR1’s role in the Il10^−/−^ model of chronic colitis. Il10^−/−^;TNFR1^fl/fl^ mice showed classic colitis features, hyperplastic crypts, increased inflammation (Fig. 2A), and elevated TNF expression compared to Il10^−/−^;TNFR1^ΔIEC^ controls (Fig. 2*B i*). Lyons et al. (6) reported that colitic expansion of the TAC compartment includes regenerative secretory progenitors with chemokine expression restricted to TACs. We observed elongation of the Reg4^+^ DCS cell zone within the TAC region in colitis (Fig. 2*B ii*), mirroring human UC. This zone was reduced in Il10^−/−^;TNFR1^ΔIEC^ mice (Fig. 2*C*). Reg4^+^ cells were enriched in Ly6a^+^ regenerating crypts of Il10^−/−^;TNFR1^fl/fl^ mice (Fig. 2*A*), suggesting a role in epithelial barrier restoration, supported by prior evidence of Reg4 as a niche factor for Lgr5^+^ stem cells (2). Similarly, Spink4^+^ DCS cells, also localized at the crypt base, were reduced in Il10^−/−^;TNFR1^ΔIEC^ mice (Fig. 2D), indicating that TNFR1 signaling drives DCS expansion in TACs during colitis. Il10^−/−^;TNFR1^ΔIEC^ mice also exhibited reduced chemokine expression, crypt hyperplasia, and histological injury (Fig. 2*E*). These findings show that epithelial TNFR1 contributes to colitis severity and Reg4^+^ DCS expansion, likely supporting epithelial repair, though additional TNFR1-independent triggers of Reg4^+^ cell expansion in inflammation cannot be excluded.

**Fig. 2.**
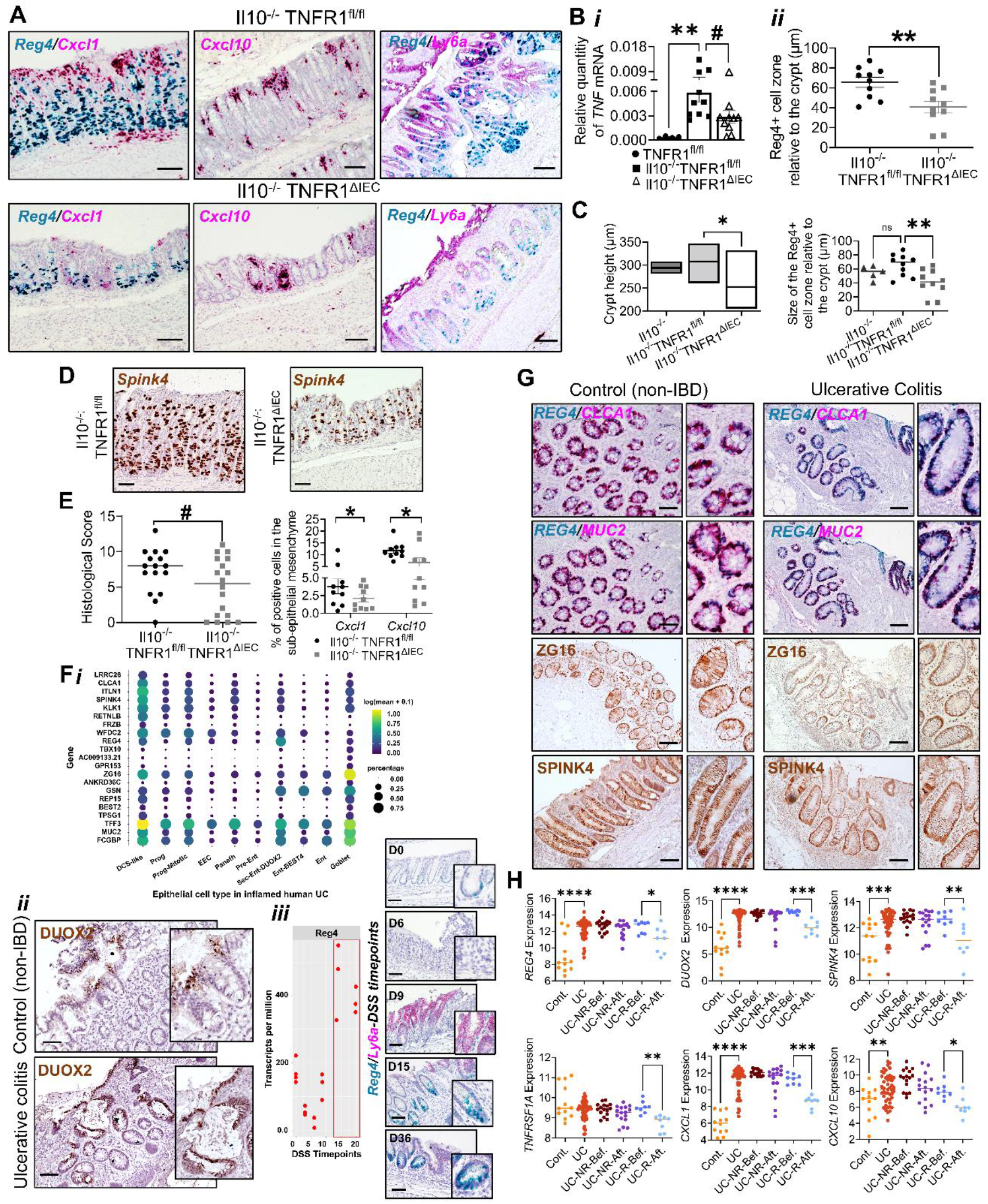
Duplex ISH for Reg4-Cxcl1, Cxcr3-Cxcl10, and Reg4-Ly6a in distal colon of *Il10*^*-/-*^*;TNFR1*^*fl/fl*^ *and Il10*^*-/-*^*;TNFR1*^*ΔIEC*^ *mice* (scale = 100 µm) (A). (i) TNF qRT-PCR; (ii) Reg4^+^ zone height relative to crypt length in *Il10*^*-/-*^*;TNFR1*^*fl/fl*^ *vs Il10*^*-/-*^*;TNFR1*^*ΔIEC*^ *mice* (B). Crypt height and Reg4^+^ zone size in Il10^-/-^, *Il10*^*-/-*^*;TNFR1*^*fl/fl*^ *and Il10*^*-/-*^*;TNFR1*^*ΔIEC*^ *mice* (n = 5-10) (C). Spink4 IHC in distal colon of *Il10*^*-/-*^ *;TNFR1*^*fl/fl*^ *and Il10*^*-/-*^*;TNFR1*^*ΔIEC*^ *mice* (scale = 100 µm) (D). Histology scores and % Cxcl1^+^/Cxcl10^+^ mesenchymal cells (n = 4-10) (E). (i) Epithelial gene expression dot plot in UC (7); (ii) DUOX2 IHC in pediatric control and UC colon (n = 5); (iii) Reg4 transcript and Reg4-Ly6a ISH in DSS-colitis (2% DSS, 5 d + 6–35 d recovery; n = 3–5) (F). REG4-CLCA1, REG4-MUC2 duplex ISH and ZG16/SPINK4 IHC in pediatric non-IBD and UC colonic crypts (n = 5; scale = 100 µm) (G). REG4, DUOX2, SPINK4, CXCL1, CXCL10 transcript levels in controls (n = 12), UC (n = 48), and infliximab-treated responders (n = 8) and non-responders (n = 16) (9) (H). Data: *t*-test or ANOVA with Tukey’s post hoc; ^#^p < 0.07, *p < 0.05, **p < 0.01, ***p < 0.001, ****p < 0.0001.

To investigate the role of Reg4^+^ cells in UC, we reanalyzed single-cell RNA-seq data from remodeled epithelial cells in the UC colon (7). REG4 was highly enriched in DUOX2^+^ secretory enterocytes co-expressing SPINK4, MUC2, ZG16, TFF3, and ITLN1, markers linked to epithelial lineage maintenance (Fig. 2*F i*). REG4, like DUOX2, localized to regenerating crypts and surface epithelium in pediatric UC patients (Fig. 2*F ii*), suggesting a role in epithelial renewal. This is supported by the DSS-colitis model, where Reg4 is specifically upregulated during recovery (days 15–35) but absent at peak injury (day 9) (8) (Fig. 2*F iii*). At baseline, REG4 marks differentiated secretory cells expressing CLCA1 and MUC2, and in UC patients, it overlaps with ZG16^+^ and SPINK4^+^ secretory cells (Fig. 2*G*, 1*B*), indicating its role in crypt recovery and epithelial barrier preservation. Finally, reanalysis of RNA-seq data from UC patients treated with anti-TNF therapy (9) shows that non-responders maintain high expression of *REG4, SPINK4, DUOX2, TNFRSF1A, CXCL1*, and *CXCL10* before and after treatment, while responders show a marked reduction (Fig. 2*H*), implicating TNF signaling in regulating these genes in UC.

## Discussion

While REG4 upregulation in UC is known (1), its regulation by TNF signaling during anti-TNF therapy remains unclear. We demonstrate that TNF induces Reg4^+^ secretory cells in both mouse and human colonic epithelium by a TNFR1-dependent mechanism. Despite species-specific localization, these cells share markers such as SPINK4 and RETNLB (11) and show conserved TNF responsiveness. We also show that pediatric and adult UC patients with primary or secondary non-response to Infliximab maintain elevated REG4, CXCL1, and CXCL10, unlike responders who normalize these markers post-treatment. Previous work suggests that factors beyond TNF, such as interferon-driven TNFR1 activation, can contextually regulate epithelial inflammation *in vivo*, reinforcing TNFR1’s role as a key regulator of REG4 even in the absence of TNF signaling (10). While sample size and model limitations prevent definitive conclusions about Reg4^+^ cell function in colitis in this study, their consistent expansion during inflammation suggests a key role in epithelial remodeling. Notably, TNF effects were observed in epithelial-only colonoids and *in vivo*, indicating an epithelial-intrinsic mechanism. Together, these findings identify a TNFR1-dependent pathway regulating Reg4^+^ cells and suggest a potential target to enhance anti-TNF therapy in UC.

## Materials and Methods

Human colonic tissue was obtained under IRB approval (CCI-13-00287, CCI-09-00093; Children’s Hospital Los Angeles) with informed consent, following the Declaration of Helsinki. Patients (mean age 8 years; 5 male, 5 female) with UC were primary or secondary non-responders to Infliximab; tissue from inflamed descending colon. Animal studies were approved by CHLA IACUC (#288). C57BL/6J Il10^−/−^ (stock #002251) and TNFR1^−/−^ (stock #002818) mice were obtained from Jackson Laboratory. TNFR1^ΔIEC^ mice were generated by crossing Villin-CreER with TNFR1^fl/fl^. Distal colonic strips were collected for RNA or organoid culture; swiss-rolled colons were fixed and sectioned at 5 μm for histology. WT mice received 2% DSS for 5 days; injury and recovery were assessed on days 6, 9, 12/15, and 35. TNFR1^+/−^ and TNFR1^−/−^ murine-colonoids were seeded at 1×10^3^ cells/well for TNF dose-response (1–100 ng/mL, 7 days). Human pediatric colonoids were treated with anti-TNFR1 or IgG isotype prior to TNF stimulation (100 ng/mL, 1 h). RNA and cDNA were isolated using Ambion^®^ PureLink^®^, Verso cDNA kits, and analyzed by qRT-PCR (TaqMan; Reg4 and Actin). RNAscope in-situ hybridization used probes for Reg4, Spink4, DUOX2, Cxcl1, Cxcl10, Cxcr3, Ly6a, CLCA1, and MUC2 with standard controls. Immunostaining employed anti-REG4, anti-ZG16, and anti-CLCA1. For RNA-Seq, day 5 organoids were harvested, sequenced (2×75 bp, HiSeq 4000), mapped (Bowtie2), quantified (RSEM), normalized (TMM), and analyzed for differential expression (edgeR, p < 0.05, >1.5-fold) with GO/pathway enrichment via ClusterProfiler. Patient transcript analyses were performed using Hegemon and GraphPad Prism. Data are available from the corresponding author; RNA-Seq data are deposited at NCBI GEO (GSE201013).

