## Supplementary Data for "TNF signaling drives expansion of Reg4+ epithelial cells in colitis"

### Supplementary Materials and Methods

*Human tissue ethical approval*

Human tissue was collected after receiving ethical approval for studies and written informed consent from patients' legal guardian, under approved Institutional Review Board CCI-13-00287 and CCI-09-00093 at Children's Hospital Los Angeles (CHLA). Human studies were conducted in accordance with the Declaration of Helsinki's criteria.

*Experimental Animals*

Animal studies were performed according to CHLA's Institutional Animal Care and Use Committee (IACUC), protocol #288 rules and regulations. $Il10^{-/-}$ (stock #002251) and $TNFR1^{-/-}$ (stock #002818) C57BL/6J-background mice were obtained from Jackson Laboratory (Maine, ME). $TNFR1^{fl/fl}$ was a kind gift from *George Kollias, Alexander Fleming Biomedical Sciences Research Center, Vari, Greece*. Intestinal epithelial-specific TNFR1 knockout mice ($TNFR1^{\Delta IEC}$) were generated by crossing Villin CreER mice with $TNFR1^{fl/fl}$ mice. $Il10^{-/-}TNFR1^{fl/fl}$ and $Il10^{-/-}TNFR1^{\Delta IEC}$ mice littermates were generated from control $Il10^{-/-}$ and $TNFR1^{fl/fl}$ and/or $TNFR1^{\Delta IEC}$ parents. Experimental and control groups, including both male and female adult mice (8-16 wks-old) were co-housed for the duration of the study.

*Mice Tissue Collection*

Dissected colons were cleared of fecal contents. Thin strips of tissue from the distal colon (2cm from rectum) were collected for RNA isolation or organoid generation. For histology, the colons were swiss-rolled, fixed in 10% formalin, and embedded in paraffin. 5 µm slices of the tissue were sectioned on the microtome.

*Murine Three-Dimensional Organoid Culture and TNF dose-response experiments*

Crypts, isolated from the distal colon of control ($TNFR1^{+/-}$), $TNFR1^{-/-}$ mice, were embedded in Matrigel® with mouse IntestiCult™ medium (STEMCELL Technologies, Vancouver, BC, no. 06005). Organoids were passaged by syringing using a 27G X 1/2 needle (Becton Dickinson, San Diego, CA, no. 305109), every 5–8 days with a 1:4 split ratio and a medium change every other day. Each experiment was repeated thrice, with >30-50 organoids/sample. Organoids were dissociated into single cells and seeded at $1x10^3$ cells/well for all experiments. Each organoid well was treated with increasing doses (1-100 ng/mL) of m-TNF (R&D systems, Minneapolis, MN; no. 410-MT) at passage for 7 days, before sample processing.

*RNAscope – In-situ hybridization*

*In-situ* hybridization (ISH) was performed according to manufacturer's recommendations for RNAscope® 2.5 HD Detection Reagents-BROWN (ACD, Newark, CA; no. 322310). The RNAscope probes included, m-*Reg4* (no. 409601), m-*Spink4* (no. 564881), h-*DUOX2* (no. 549111), h-*REG4* (no. 312071), m-*Cxcl1* (no. 407721-C2), m-*Cxcl10* (no. 408921-C2), m-*Cxcr3* (no. 402511-C2), m-*Ly6a* (no. 427571-C2), h-*CLCA1* (no. 483161-C2), and h-*MUC2* (no. 312879-C2). RNAscope® Control Slide-Mouse 3T3 Cell Pellet (no. 310023) and RNAscope® Control Slide -Human Hela Cell Pellet (no. 310045) were included as controls.

*Histochemistry*

Immunostaining was performed according to standard protocols. Antibodies used in this study included anti-REG4 (Sigma Aldrich, St. Louis, MO, no. HPA046555), anti-ZG16 (Sigma Aldrich, no. HPA052512), and anti-CLCA1 (Sigma Aldrich, no. HPA052787).

*RNA Isolation, cDNA Synthesis and qRT-PCR*

Total RNA from colon or organoids was extracted using the Ambion® PureLink® RNA Isolation Kit (Thermofisher Scientific, no. 12183025). cDNA was synthesized from 50-300 µg of treated RNA for each sample using Verso cDNA Synthesis Kit (Thermofisher Scientific, no. AB1453B) using a CFX384 Touch™ Real-Time PCR Detection System (Bio-Rad, Irvine, CA). qRT-PCR for a subset of primers was performed using the TaqMan Fast Advanced Master Mix (ThermoFisher Scientific, no. 4444557) on a Bio-Rad iQ5 thermocycler. Primer and probe oligonucleotides were obtained from Integrated DNA Technologies (Newark, NJ). *Reg4* (Mm.PT.58.12037171) and *Actin* (Mm.PT.39a.22214843). qRT-PCR reactions contained 5.2 µL SYBR® Premix Ex Taq™ II (Tli RNase H Plus), ROX Plus (Takara, Mountain View, CA; no. RR82WR), 0.2 µL each of sense and antisense primers, h-*β-actin*: 5'-3'- *AGCACGGCATCGTCACCAACT*; 3'-5'- *TGGCTGGGGTGTTGAAGGTCT* and h-*Reg4*: 5'-3'- *GCCCCGCCATCCCTT*; 3'-5'- *CTGCTCGAGACAGCCAGAGA*, at 10 µM, 2 µL cDNA or no reaction control with 2.4 µL nuclease free water to a final volume of 10 µL. Samples were heated to 95ºC for 3 min followed by 37 cycles of 94ºC for 15 s, 54ºC for 20 s, and 72ºC for 25 s. Data were analyzed using the comparative ΔΔCt method.

*Human organoid culture, TNF treatment and anti-TNFR1 experiments*

Colonoids were generated from the descending colon of control pediatric (non-IBD) patients using pre-established techniques. Colonoids were passaged by syringing using a 27G X 1/2 needle in a 1:4-6 split ratio, every 8-10 days. Medium was changed every 2-3 days. All experiments were carried out from passage 3-8. H-organoids at day 10 were pre-treated with 0.5 µg/mL anti-TNFR1 and its IgG isotype control (Sino Biological, PA, no. 150496-M08H) in parallel for 1h before passage and then seeded at $1x10^3$ cells/well, for a 10-day exposure. On day 10 the control untreated colonoids and those cultured with anti-TNFR1, and IgG isotype were treated with 100ng/mL TNF (R&D systems, no. 210-TA-100) for 1 h before sample processing.

*RNA-Seq*

Total RNA from day 5 control and $TNFR1^{-/-}$ primary organoids (passage 3) was extracted using the Ambion® PureLink® RNA Isolation Kit. mRNA transcripts were prepared for 2x75 bp sequencing on a HiSeq 4000 (Illumina). Library preparation and sequencing were performed by CHLA's Single Cell, Sequencing, and CyTOF Core Laboratory (Los Angeles, CA). The reads were first mapped to the latest UCSC transcript set using Bowtie2 version 2.1.0 and the gene expression level was estimated using RSEM v1.2.15. TMM method was used for the normalization of the raw count. Differentially expressed genes were identified using the edgeR program. Genes showing altered expression with $p < 0.05$ and more than 1.5-fold changes were considered differentially expressed. Cluster profiler was used for the GO and pathway enrichment analysis.

*DSS Colitis*

C57BL/6J-mice were given 2% DSS (36–50 kDa molar mass; MP Biomedicals) in distilled water for 5 days. Injury and recovery timepoints were collected on days 6, 9, 12/15 and 35. Whole colonic mucosal cells were examined for single cell RNA sequencing analysis[9] and tissue was collected for histology.

*Bioinformatics Analysis of Gene-Expression Array Databases*

Analysis of target genes expression was determined from datasets generated by *Arijs et al, 2009* (4), using Hegemon (10). Raw data for each target from non-IBD controls and UC patients before or after treatment with Infliximab was imported into the GraphPad Prism Software. Data were analyzed by a one-way Anova and Šidák's Posthoc test, where *$p < 0.05$, **$p < 0.01$, ***$p < 0.001$, and ****$p < 0.0001$.

Data, analytic methods, and study materials will be made available from the corresponding author on reasonable request. The RNA-Seq dataset is available at NCBI Gene Expression Omnibus, identified by the accession number GSE201013.
